# OPUS-Mut: studying the effect of protein mutation through side-chain modeling

**DOI:** 10.1101/2022.05.10.491420

**Authors:** Gang Xu, Qinghua Wang, Jianpeng Ma

## Abstract

Predicting the effect of protein mutation is crucial in many applications such as protein design, protein evolution, and genetic disease analysis. Structurally, the mutation is basically the replacement of the side chain of a particular residue. Therefore, accurate side-chain modeling is useful in studying the effect of mutation. Here, we propose a computational method, namely OPUS-Mut, which significantly outperforms other backbone-dependent side-chain modeling methods including our previous method OPUS-Rota4. We evaluate OPUS-Mut by four case studies on Myoglobin, p53, HIV-1 protease, and T4 lysozyme. The results show that the predicted structures of side chains of different mutants are consistent well with their experimentally determined results. In addition, when the residues with significant structural shifts upon the mutation are considered, it is found that the extent of the predicted structural shift of these affected residues can be correlated reasonably well with the functional changes of the mutant measured by experiments. OPUS-Mut can also help one to identify the harmful and benign mutations, and thus may guide the construction of a protein with relatively low sequence homology but with similar structure.

## Introduction

Protein amino-acid mutations play a key role in protein engineering and protein evolution (1), which are also the root causes of many genetic diseases (2). Therefore, it is crucial to predict the change of mutant properties against their original unmutated counterparts (wild-type). In recent years, with the development of deep learning techniques, many successful researches have been performed to predict the change of different properties for mutation, such as thermodynamic stability (3, 4), enzyme activity (5), and loss- or gain-of-function (6, 7). However, limitations still exist. For example, for protein stability prediction, due to the sequence similarity between proteins used in training and test datasets, the methods tend to overestimate the prediction performance (8).

From a structural point of view, a mutation is basically the replacement of the side chain of the corresponding residue. In this case, we study the effect of protein mutation through side-chain modeling. By comparing the differences between the predicted unmutated (wild-type) side chains and the predicted mutant side chains, we can infer the extent of structural perturbation and the affected residues whose predicted side chains are significantly shifted due to the mutation. To quantify the extent of structural perturbation, we use the summation of the differences of all predicted side-chain dihedral angles (from χ_1_ to χ_4_) over all residues between the wild-type and mutant. In this paper, for the purpose of clarity, we name this summation of differences over all residues as *S_diff_*. It is assumed that larger *S_diff_* corresponds to more severely predicted structural perturbation, which means the mutation is more likely to be harmful. From the affected residues, we can correlate them with the functional changes. In protein engineering, for example, with the possibility of inferring the affected residues for a particular mutation, we may selectively avoid the type of mutations that could potentially impair the functions related to certain residues.

There are some advantages in studying the effect of mutation through a general side-chain modeling method, in oppose to computational methods using the experimentally obtained mutational-functional changes data. They are 1) it has no system dependence and can be generalized to any target without requiring further system-specific data for fine-tuning; 2) it focuses on the side-chain modeling ability and is not limited by the shortage of the experimental data of mutant properties; 3) it is straightforward to interpret since it is based on the predicted structural shift.

In recent years, many protein structure prediction methods have been proposed (9–12). Among them, AlphaFold2 (11) is the best and, in some cases it delivers the predictions close to the experimental results. However, AlphaFold2 may not be suitable for the protein mutation task (13). Firstly, they are insensitive to the single site mutation. In our study, we find that the predicted backbone for the wild-type sequence and the sequence with single site mutation are usually identical, maybe because they all depend on the results of multiple sequence alignment, which are similar to each other when the query sequences are almost the same. In contrast, OPUS-Mut adopts deep neural network to capture the local environment for each residue, therefore, it is sensitive to the point mutation. Secondly, the structure prediction methods, like that of AlphaFold2, usually deliver predictions carrying stochastic noise. Sometimes, the structural difference between their predicted structures is larger than the difference caused by the mutation. Therefore, it may be difficult to distinguish whether the structural difference is caused by the mutation or is merely the prediction noise. In this case, using the side-chain modeling results with a fixed backbone (here, we use wild-type experimental backbone for both wild-type and mutant cases) as a first order approximation is a rational trade-off for studying protein point mutation from structural perspective.

For backbone-dependent side-chain modeling, many successful methods have been developed. For sampling-based methods (14–17), they sample the rotamers from the rotamer library, and use their scoring functions to determine the best rotamer for each residue with the minimal score. However, this kind of methods is limited by the discrete rotamers in the rotamer library and the accuracy of the scoring function. Recently, some deep learning-based methods have been proposed (18, 19), which improve the accuracy of side-chain modeling by a large margin.

In this research, we propose a computational method, named OPUS-Mut, which is mainly based on our previous work OPUS-Rota4 (19), but with some improvements. In OPUS-Mut, we modify our loss function for the predicted dihedral angles according to that in AlphaFold2 (11). To increase the sensitivity of OPUS-Mut towards the point mutation, we add an additional neural network node to predict the root-mean-square-deviation (RMSD) of the predicted side chain against its native counterpart for each residue. Therefore, OPUS-Mut can evaluate the correctness of its prediction at each site. In our study, OPUS-Mut outperforms OPUS-Rota4 and other backbone-dependent side-chain modeling methods on three native backbone datasets. The excellent side-chain modeling ability of OPUS-Mut is the foundation of this research.

For further evaluation, four case studies (Myoglobin, p53, HIV-1 protease and T4 lysozyme) have been conducted, from either structural or functional perspectives. From structural perspective, we adopt the mutants that have experimentally determined structures, and focus on the accuracy of the predicted side chains comparing to their experimental counterparts. From functional perspective, we focus on the extent of structural perturbation and the affected residues whose predicted side chains are significantly shifted due to the mutation, and the correlation between them and the functional changes. The results show that OPUS-Mut can be used in the following cases: 1) it can be used to infer the functional changes of the mutation from structural perspective; 2) it can be used to infer the affected residues due to the mutation, therefore avoid the unwanted effect on specific sites in the case of protein engineering; 3) it can help us to identify the harmful and benign mutations. As one of the applications, it may guide us to construct a relatively low-homology mutant sequence but with similar structure, thus may be helpful in studying protein engineering and protein evolution.

## Results

### Side-chain modeling performance of OPUS-Mut

We compare OPUS-Mut with two sampling-based methods SCWRL4 (16) and OSCAR-star (17), and two deep-learning based methods DLPacker (18) and OPUS-Rota4 (19), on three native backbone test sets CAMEO60, CASP14 and CAMEO65. In terms of the percentage of correct prediction with a tolerance criterion 20° for all side-chain dihedral angles (from χ_1_ to χ_4_), OPUS-Mut outperforms other methods either measured by all residues (Figure 1) or measured by core residues only (SI appendix, Fig. S1). The percentage of correct prediction for each type of residue on CAMEO65 is also listed in SI appendix, Table S1 (all residues) and SI appendix, Table S2 (core residues).

**Figure 1.**
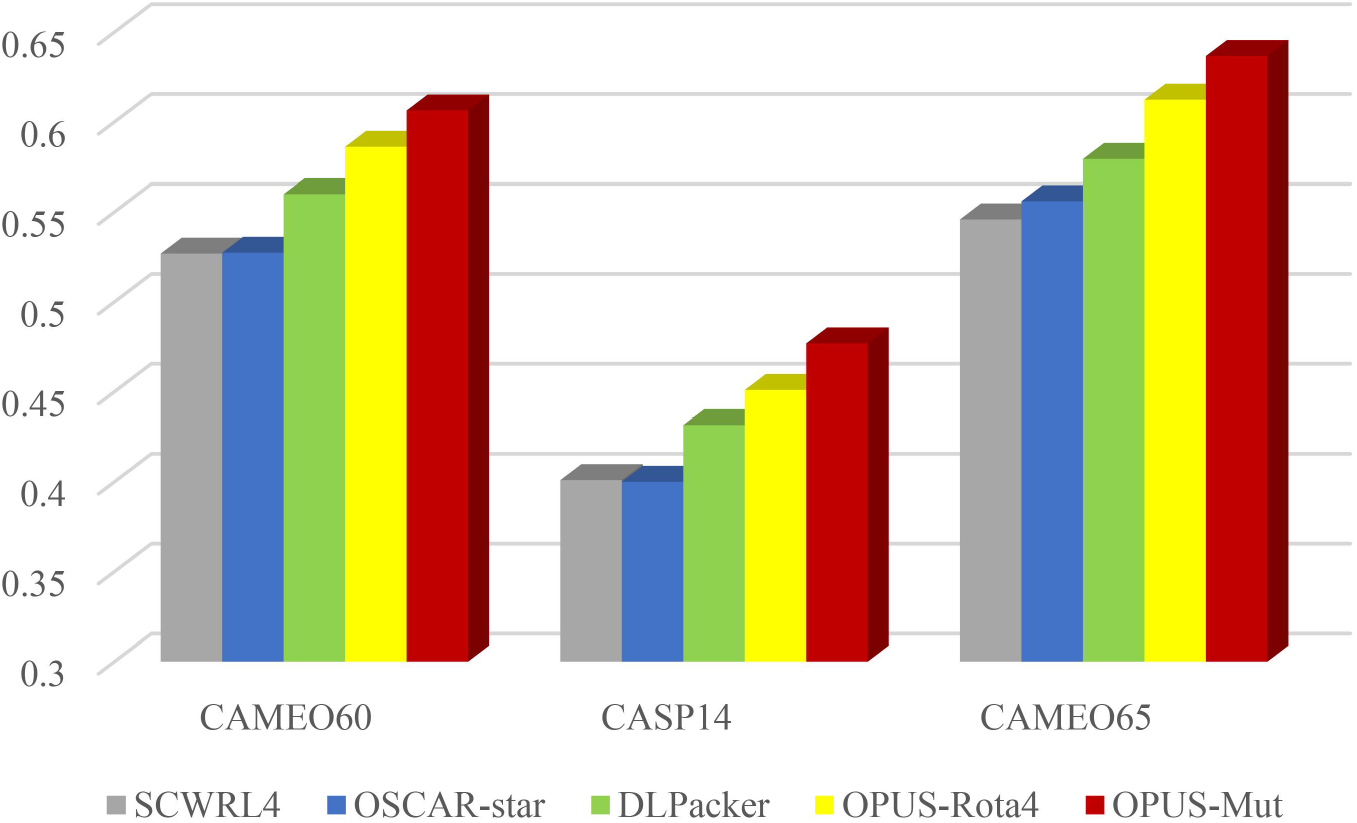
The percentage of correct prediction with a tolerance criterion 20° for all side-chain dihedral angles (from χ_1_ to χ_4_) of different methods measured by all residues.

### Case study: Myoglobin

Myoglobin is an essential hemoprotein that regulates cellular oxygen concentration in striated muscle. It can bind oxygen reversibly by its heme prosthetic group (20).

#### From structural perspective

Previous study showed that the substitutions (V68A, V68I, V68L and V68F) at the position 68 do not affect the secondary or tertiary structure of the protein significantly, and the largest changes caused by these substitutions are the distortions of the interior spaces (21). Meanwhile, the structure of these mutants (V68A (PDB: 1MLF), V68I (PDB: 1MLM), V68L (PDB: 1MLQ) and V68F (PDB: 1MLJ)) have been determined by Quillin *et al*. (21). Therefore, examining the side-chain modeling results for these four substitutions is an ideal way to verify the sensitivity of OPUS-Mut towards the single-site mutation.

Following the research from Nienhaus *et al*. (22), we download the structure of the wild-type (PDB: 2MGK) sperm whale MbCO (23). Then, we replace the Val at position 68 by Ala, Ile, Leu and Phe, and reconstruct their side chains with OPUS-Mut, respectively.

As shown in Figure 2, the side-chain modeling results for each mutant are consistent well with their X-ray crystallographic results (21). From the side-chain prediction results of mutant V68F (Figure 2f), we can infer that the benzyl side chain that partially fills the Xe4 cavity (region C in Figure 2a) may become a steric barrier to the ligand association. This assumption has been confirmed through experimental approaches (21, 22).

**Figure 2.**
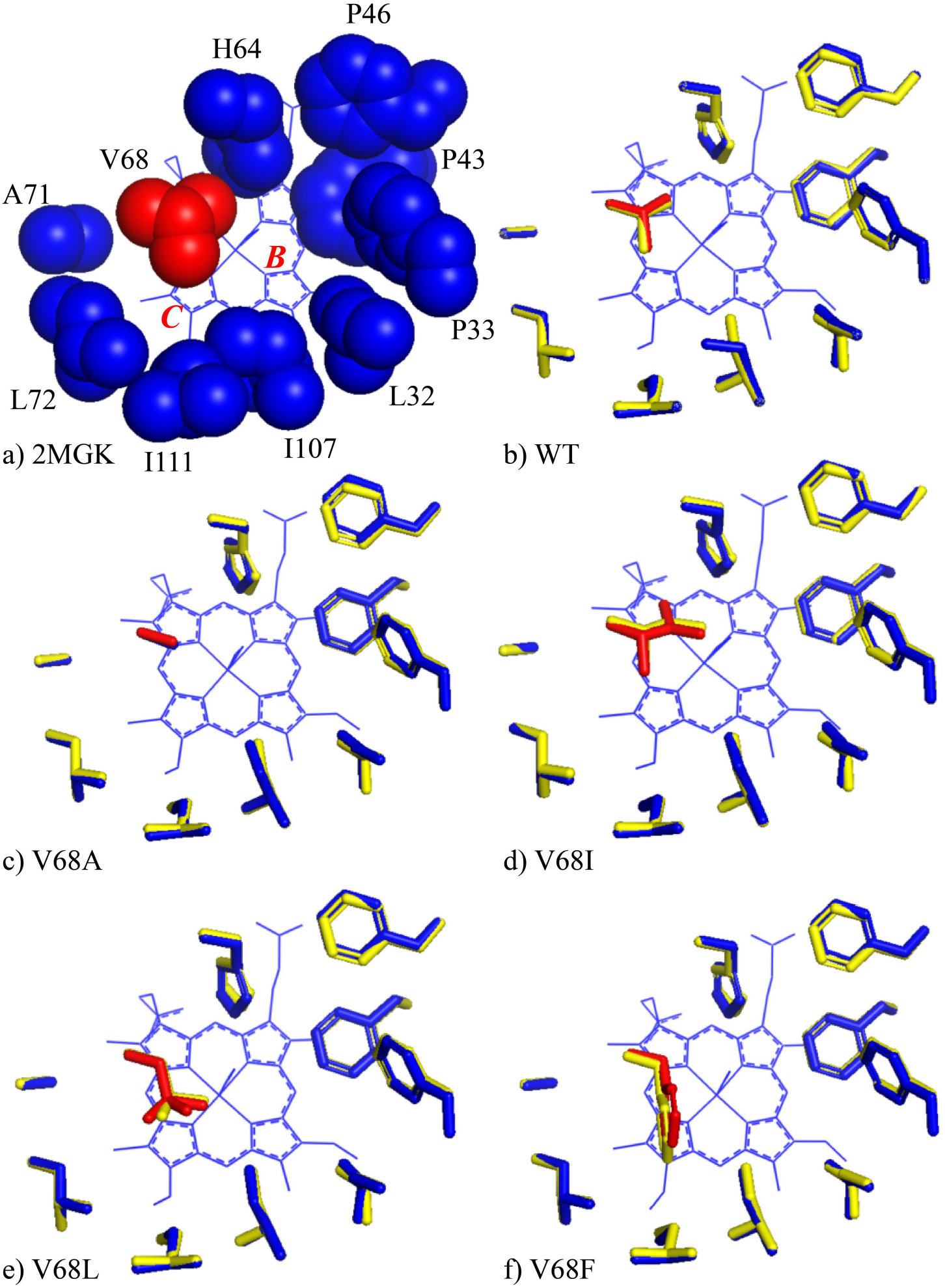
Side-chain modeling results of wild-type MbCO and its mutants. a) The local environment around V68 in the wild-type MbCO crystal structure (PDB: 2MGK). The Xe4 pocket consists of region B and C. b) The side-chain structures in the wild-type structure (PDB: 2MGK) and the side-chains predicted by OPUS-Mut. c)-f) The crystal and predicted side-chain structures for each mutant: c) the V68A mutant (PDB: 1MLF); d) the V68I mutant (PDB: 1MLM); e) the V68L mutant (PDB: 1MLQ); and f) the V68F mutant (PDB: 1MLJ). The crystal side-chain structure at the mutation site is marked in red, the crystal side-chain structures of the nearby residues are marked in blue, and the predicted side chains are marked in yellow. The results indicate that the predicted mutant side chains (yellow) are consistent with their crystal structures (red and blue). Note in the side-chain modeling calculation, the heme group is not included. However, for the purpose of illustration, the heme group is shown in each panel.

### Case study: p53

The tumor suppressor p53 plays an important role in maintaining the genomic integrity of the cell, and it is mutated in half of all human cancers (2). In the last few decades, many researches have been done to study the different properties of its mutations (24–31).

#### From structural perspective

Fersht group constructed *T*-p53C, a superstable quadruple mutant (M133L/V203A/N239Y/N268D) of human p53 core domain (30) and determined its structure (PDB: 1UOL) (29). Furthermore, based on *T*-p53C, the experimental structure of a series of mutants were determined (27, 28), including R273C (PDB: 2J20), R273H (PDB: 2BIM), F270L (PDB: 2J1Z), V143A (PDB: 2J1W), Y220C (PDB: 2J1X), R282W (PDB: 2J21), and triple mutant T123A/H168R/R249S (PDB: 2BIQ).

In this study, we download the *T*-p53C (PDB: 1UOL chain A). Then, we substitute the residues according to the corresponding mutations mentioned above and reconstruct their side chains with OPUS-Mut, respectively.

As shown in Figure 3 and SI appendix, Fig. S2, our side-chain modeling results for each mutant are very close to their experimental counterparts, especially for the mutations inside the protein: R273C (Figure 3a), R273H (Figure 3b), F270L (Figure 3c) and V143A (Figure 3d). For the mutation on the loop area Y220C (SI appendix, Fig. S2b), the prediction is also accurate and the biases are mainly caused by the main-chain perturbation, which means the main chains of the loop area in the original structure (PDB: 1UOL chain A) we used for side-chain modeling is not strictly identical to that in the mutant structure (PDB: 2J1X). For R282W (SI appendix, Fig. S2c), the side-chain predictions for its nearby residues that are close to the deoxyribonucleic acid are less accurate than the others, because OPUS-Mut doesn’t take the effect of deoxyribonucleic acid into account. The results for T123A/H168R/R249S (SI appendix, Fig. S2d) indicate the good performance of OPUS-Mut in modeling the side chains of multiple mutant.

**Figure 3.**
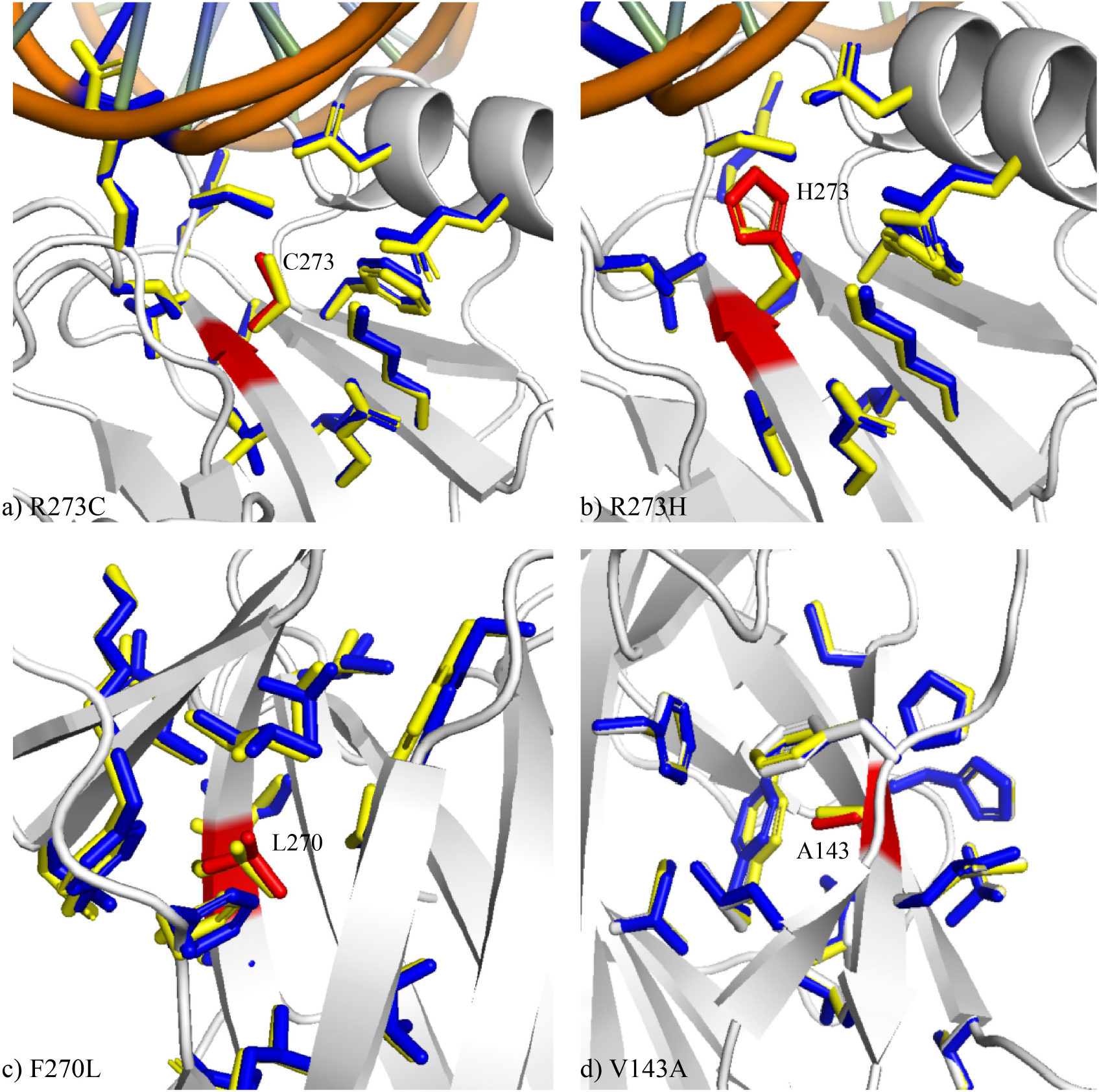
Side-chain modeling results of *T*-p53C mutants. a) The R273C mutant (PDB: 2J20); b) the R273H mutant (PDB: 2BIM); c) the F270L mutant (PDB: 2J1Z); and d) the V143A mutant (PDB: 2J1W). In all panels, the side chains from the crystal structures are marked in red for the mutation sites, and in blue for neighboring residues within 5 Å from the mutation sites. The side chains predicted by OPUS-Mut are marked in yellow.

#### From functional perspective

In this section, we focus on three Zn^2+^ region mutations (R175H, M237I and C242S) and a DNA region mutation (R282W) (2). We download the p53-DNA complex (PDB: 1TUP chain B) (32) and substitute the residues according to the mutations mentioned above. Then, we reconstruct their side chains with OPUS-Mut respectively. Note that, according to Bullock *et al*., Zinc-binding site consists of four residues: C176, H179, C238 and C242 (2).

As shown in Figure 4, all the affected residues are shown and marked in green. Apart from the affected residues, the side chains of other residues in the protein remain relatively unshifted. Figure 4a-c show that the residues at Zinc-binding site (C238 and C242 in Figure 4a, C242 in Figure 4b, and H179 in Figure 4c) are the affected residues in each mutant, which indicate the Zn^2+^ region may be modified in these three mutants.

**Figure 4.**
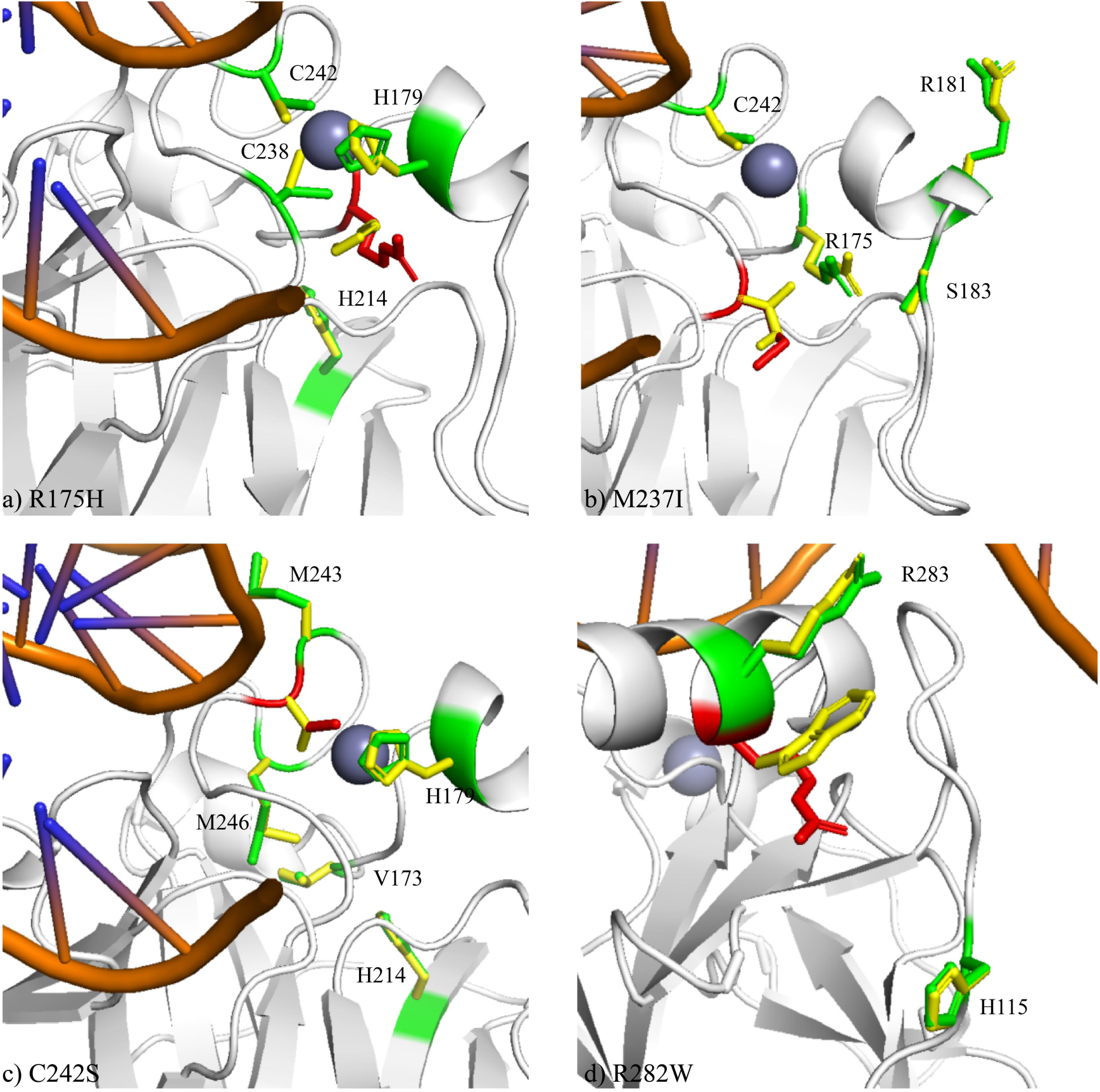
Side-chain modeling results of p53-DNA complex mutations. a) The R175H mutant; b) the M237I mutant; c) the C242S mutant; and d) the R282W mutant. The mutation site is marked in red, and all the affected residues are shown and marked in green. The affected residue refers to the residue whose mean absolute error of all predicted side-chain dihedral angles between the wild-type and mutant is greater than 5 degree. The predicted side chains of the wild-type (PDB: 1TUP) are marked in green (red at the mutation site), and the predicted side chains of the mutants are marked in yellow.

For R282W (Figure 4d), R283 and H115 are its affected residues. According to Rippin *et al*., R283 belongs to a DNA binding determinant (31). We assume that this may be a clue for R282W’s binding affinity loss. The predicted side-chain conformation changes at H115 maybe a prediction bias since it’s far away from W282.

### Case study: HIV-1 Protease

HIV-1 protease is an important anti-AIDS drug target (33), and many studies have been conducted to investigate its different properties (33–40).

#### From structural perspective

Ala *et al*. determined the structure of the mutant HIV-1 proteases (V82F, I84V) complexed with cyclic urea inhibitors (35). Since the backbone structure of HIV-1 protease may be shifted along with different inhibitors. In this case study, instead of using the wild-type backbone structure and replace the mutation residues at the corresponding sites, we download the mutant backbone structure (PDB: 1MEU chain A) directly and reconstruct its side chains.

As shown in Figure 5, for mutation residues F82 and V84, the side-chain modeling results are very close to the experimental results. The predictions for their nearby residues (< 5 Å) are also accurate, except the predictions for the surface residues R8 and E21.

**Figure 5.**
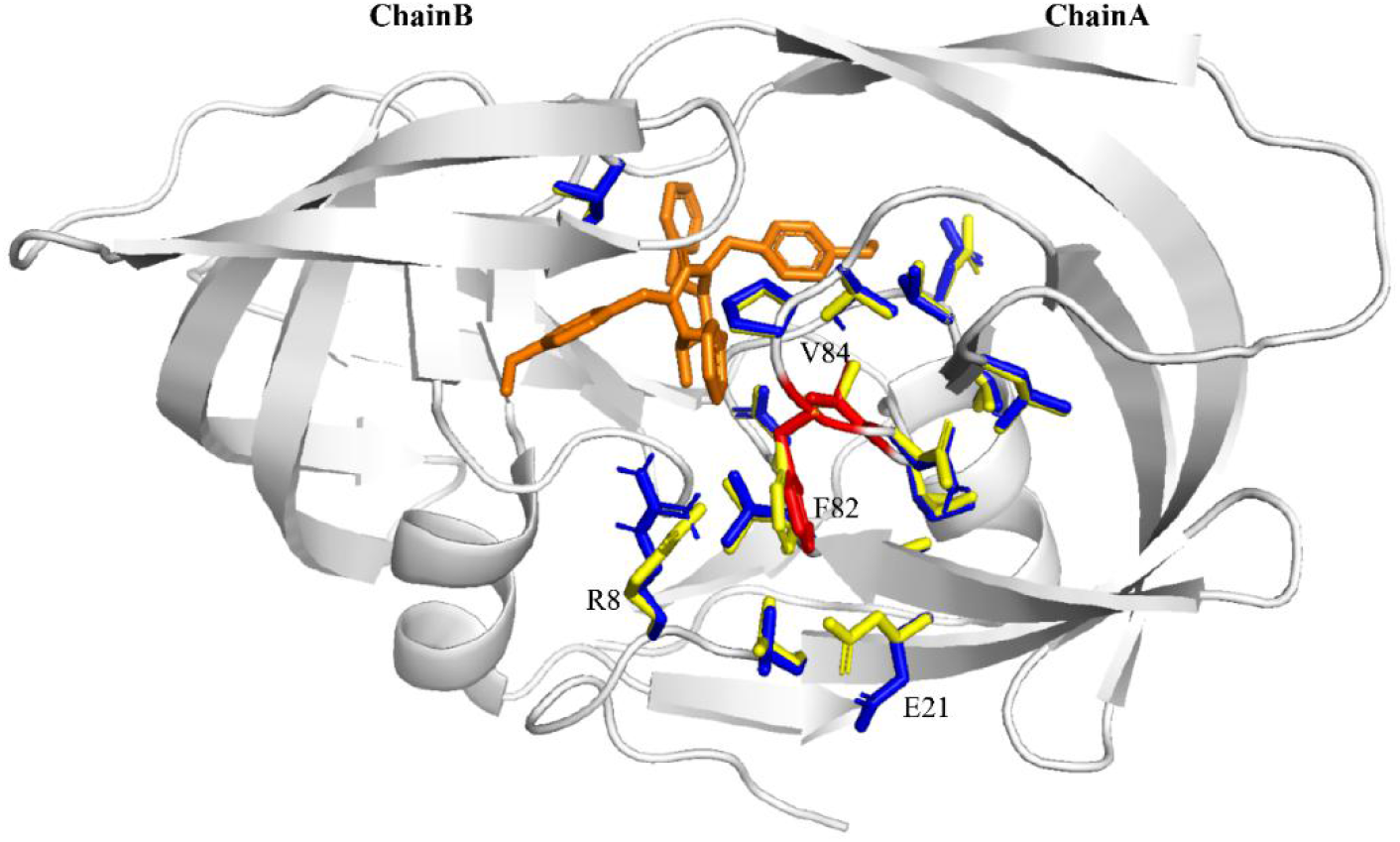
Side-chain modeling results of HIV-1 proteases complexed with cyclic urea inhibitors. The side chains from the crystal structure (PDB: 1MEU) are marked in red for the mutation sites (V82F, I84V), and in blue for neighboring residues within 5 Å from the mutation sites. The cyclic urea inhibitor is shown in brown. The side chains predicted by OPUS-Mut are marked in yellow.

#### From functional perspective

The complete mutagenesis of HIV-1 protease has been performed by Loeb *et al* (40). In that study, they measured 336 single missense mutations and their corresponding phenotypes. The phenotypes were categorized into three groups based on their ability to process the Pol precursor in *E. coli:* negative (no mature processed products), intermediate (some mature processed products), and positive (processed similarly to the wild-type). In this case study, we collect their results to form a HIV-1 protease mutagenesis dataset that contains 319 results (because some results are hard to distinguish from their paper, especially for upper-case *I* and lower-case *l*). The dataset we collected contains 153 negative phenotype data, 66 intermediate phenotype data, and 100 positive phenotype data (SI appendix, Table S3).

Following the research from Masso *et al*. (5), we download the structure of the wild-type HIV-1 protease with 99 residues in length (PDB: 3PHV) (41). We predict the side chains for all 99 × 19 = 1881 possible single site mutations. For each mutation, we sum up the differences of all side-chain dihedral angles (from χ_1_ to χ_4_) between the predicted wild-type side chains and the predicted mutant side chains, and use it as an indicator for the extent of structural perturbation due to the mutation. Here, beside the summation of differences over all residues *S_diff_*, we use *S_diff_critical_* to denotes the summation of differences on some critical residues (such as the residues in active site).

In all 1881 possible single site mutations, only 319 of them have experimental results (the data in the HIV-1 protease mutagenesis dataset), therefore, we use them to do the statistical analysis. As shown in Figure 6a, the mean of *S_diff_* for the mutations in negative (0), intermediate (1) and positive (2) phenotype group is 28.3, 23.0 and 21.7, respectively. The median of them is 27.0, 20.5 and 17.0, respectively. Both mean and median of these three groups show that more severely predicted structural perturbation corresponds to a higher probability of loss-of-function. Among 319 mutations, 7 out of the top 10 mutations with the highest *S_diff_* belong to the negative phenotype group, 2 out of the top 10 mutations belong to the intermediate phenotype group, and only 1 out of the top 10 mutations belongs to the positive phenotype group.

**Figure 6.**
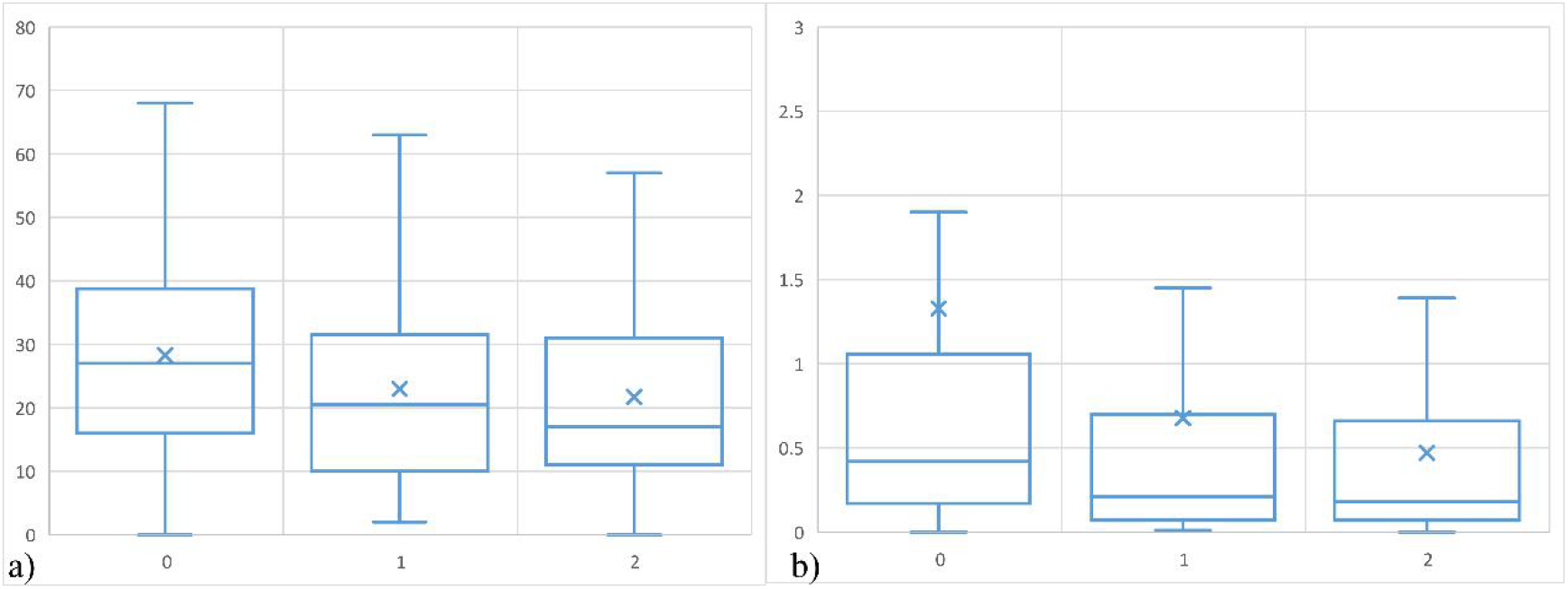
The summation of differences of all side-chain dihedral angles (from χ_1_ to χ_4_), either summed over all residues (*S_diff_*) or summed over three active site residues (*S_diff_critical_*), between the predicted wild-type side chains and the predicted mutant side chains on HIV-1 protease mutagenesis dataset. In the Box plot, “x” represents the mean of each group, the line inside the box represents the median of each group. The data from three groups (negative (0), intermediate (1) and positive (2) phenotype group) are shown. The mutations on the first and the last residues (P1 and F99) are ignored because the side-chain modeling may have a bias on the end of the sequence. a) shows the differences (*S_diff_*) summed over all residues. To avoid the influence of outliers, if the difference between the two residues is smaller than 1 degree, we set it to be 0; if it’s smaller than 5, we set it to be 1; if it’s smaller than 10, we set it to be 2; if it’s smaller than 20, we set it to be 3; if it’s larger than 20, we set it to be 4. b) shows the differences (*S_diff_critical_*) summed over three active site residues (D25, T26 and G27).

According to Mager *et al*. (34), D25, T26 and G27 are the key residues involved in the active site of HIV-1 protease. To find out the impact of each mutant on the active site, we sum up the differences (*S_diff_critical_*) on these three residues exclusively (Figure 6b). The mean of them is 1.33, 0.68 and 0.47, respectively. The median of them is 0.42, 0.21 and 0.18, respectively. Comparing to the three groups obtained by summing up the differences (*S_diff_*) over all residues, the groups obtained by summing up the differences (*S_diff_critical_*) on three active site residues are more distinguishable. Among 319 mutations, 7 out of the top 10 mutations with the highest *S_diff_critical_* belong to the negative phenotype group, 2 out of the top 10 mutations belong to the intermediate phenotype group, and only 1 out of the top 10 mutations belong to the positive phenotype group.

In Table 1, we list the top 10 mutations with the highest *S_diff_critical_* on three active site residues (D25, T26 and G27) in all 1881 possible single site mutations. The mutations on residues 24-28 are ignored since they are too close to the active site. Among these predicted top 10 mutations, V82I (39) and V82E (42) have been confirmed that they can reduce the inhibitor affinity. In addition, V82, I84 and L90 are the hot spots of the drug resistance mutation (43). Therefore, OPUS-Mut would be a useful tool in finding the possible harmful mutations based on their influences on the protein’s critical region.

**Table 1.** The top 10 HIV-1 protease mutations with the highest *S_diff_critical_* on three active site residues (D25, T26 and G27) in all 1881 possible single-site mutations.

| Mutant | $S_{diff\_critical}$ |
| --- | --- |
| P9Y | 7.32 |
| V82I | 7.17 |
| V82G | 7.10 |
| I84N | 6.76 |
| L90R | 6.63 |
| P9F | 6.31 |
| I84D | 5.69 |
| L90Y | 5.68 |
| A22M | 5.22 |
| V82E | 5.01 |

For each of 1881 possible single site mutations, we sum up the differences (*S_diff_*) of all side-chain dihedral angles between the predicted wild-type side chains and the predicted mutant side chains over all residues (with the differences on three active site residues multiplied by a factor of 10 in order to emphasize the significance of the active site residues). Then, for each residue site, among its 19 possible mutations, we keep the one with the lowest *S_diff_*. Therefore, we get 99 sequences, each with a single site mutation, that have the smallest structural perturbation. We sort 99 sequences according the value of their *S_diff_*. For a desired mutation rate, for example, 10% in Type 1, Table 2, we combine the top 10 sequences with the lowest *S_diff_* to construct a sequence with 10% of mutation rate. For comparison, another type of multiple mutation sequence is constructed reversely, by combining the sequences with the highest *S_diff_* in the procedure mentioned above (Type 2, Table 2). We use AlphaFold2 (11) without template to predict the structure for the sequences with 10%, 20%, 30%, 40%, and 50% of mutation rate. As shown in Table 2, by preferably using the minimally disturbing mutations (with the lowest *S_diff_*), the constructed multiple mutation sequences (Type 1, Table 2) tend to cause smaller structural perturbation than those (Type 2, Table 2) constructed by preferably using the maximally disturbing mutations (with the highest *S_diff_*). For example, the TM-score of the Type 1 sequence with 50% of mutation rate is 0.85, which is still reasonably high (note the TM-score of the wild-type structure prediction is 0.93). The details of the structure and sequence of this multiple mutant are shown in SI appendix, Fig. S3.

**Table 2.** The TM-score of each HIV-1 protease (with 99 residues in length) multiple mutation sequence predicted by AlphaFold2. “Type 1” represents the TM-score of the sequences with 10%, 20%, 30%, 40%, and 50% of mutation rate, constructed by preferably using the minimally disturbing mutations with the lowest *S_diff_*. “Type 2” represents the TM-score of the multiple mutation sequences constructed by preferably using the maximally disturbing mutations with the highest *S_diff_*.

|  | Wild-type | 10% | 20% | 30% | 40% | 50% |
| --- | --- | --- | --- | --- | --- | --- |
| Type 1 | 0.93 | 0.91 | 0.90 | 0.86 | 0.85 | 0.85 |
| Type 2 | - | 0.83 | 0.82 | 0.82 | 0.62 | 0.50 |

### Case study: T4 lysozyme

T4 lysozyme is a monomeric protein which contains 164 residues that hydrolyzes peptidoglycan (44). Many studies have been performed to investigate the different properties of its mutations, especially by Matthews’s group (45–50).

In this case study, we download the structure of the wild-type T4 lysozyme (PDB: 2LZM) (45), and predict the side chains for all 164 × 19 = 3116 possible single site mutations. According to the PDB file of 2LZM (45), E11 and D20 are the active site residues, L32, F104, S117 and N132 are the binding site residues.

#### From functional perspective (protein stability)

Masso *et al*. (51) collected the experimental stability changes of 293 T4 lysozyme mutants at physiological pH from ProTherm (52). From the figure of the mutational array in their paper (51), we identify 292 of them, in which 80 of them will increase the stability and 212 of them will decrease the stability (SI appendix, Table S4).

As shown in SI appendix, Fig. S4a, the mean of *S_diff_* for the mutations in decreased stability group and increased stability group is 13.2 and 10.0, respectively. The median of them is 10.0 and 7.0, respectively. Among 292 mutations, 9 out of the top 10 with the highest *S_diff_* belong to the decreased stability group. The results are consistent with our assumption that more severely predicted structural perturbation corresponds to a higher probability of stability decrease.

We also sum up the differences (*S_diff_critical_*) of all side-chain dihedral angles between the predicted wild-type side chains and the predicted mutant side chains on the binding site residues (L32, F104, S117 and N132) of T4 lysozyme exclusively (SI appendix, Fig. S4b). The mean of them is 26.6 and 14.0, respectively. The median of them is 3.1 and 1.7, respectively. Among 292 mutations, all of the top 10 with the highest *S_diff_critical_* belong to the decreased stability group. The results also indicate the importance of critical residues on distinguishing different type of mutations.

Same as the study on HIV-1 protease, for all 3116 possible single site mutations, we sum up the differences (*S_diff_*) of all side-chain dihedral angles between the predicted wild-type side chains and the predicted mutant side chains on all residues, with the differences on two active site residues and four binding site residues multiplied by a factor of 10. Two types of sequences with 10%, 20%, 30%, 40%, and 50% of mutation rate are constructed: 1) the first type of multiple mutation sequences is constructed by preferably using the minimally disturbing mutations (with the lowest *S_diff_*), 2) the second type of multiple mutation sequences is constructed by preferably using the maximally disturbing mutations (with the highest *S_diff_*). The results also indicate that the combination of the minimally disturbing mutations may lead to a multiple mutation sequence with smaller structural perturbation (SI appendix, Table S5).

#### From functional perspective (enzyme activity)

Masso *et al*. (5) collected the experimental enzyme activities of 2015 T4 lysozyme mutants obtained by Rennell *et al*. (53). Among them, 1552 mutants are labeled as active, 263 mutants are labeled as inactive.

In all 3116 possible single site mutations, the top 10 mutations with the highest *S_diff_critical_* on two active site residues (E11 and D20) are as following: 2 of them have no experimental data, 7 of them belong to the inactive group in the experimental enzyme activity data, and only one of them belongs to the active group. For further illustration, in Table 3, we list the top 20 mutations under the following rules: firstly, we exclude the mutations without experimental data; secondly, for a residue site with many mutations existed, we show up to 3 mutations (such as the mutations in Q105 in Table 3). The results show that 15 out of the top 20 mutations belong to the inactive group.

**Table 3.** The selected top 20 T4 lysozyme mutations with the highest *S_diff_critical_* on two active site residues in all 3116 possible single site mutations. ‘Rank’ refers to the real rank before selection. ‘Activity’ refers to the experimental enzyme activity, ‘N’ denotes inactive, ‘P’ denotes active.

| Rank | Mutant | $S_{diff\_critical}$ | Activity |
| --- | --- | --- | --- |
| 2 | T26Y | 15.68 | N |
| 3 | T142E | 15.6 | N |
| 4 | T26R | 15.08 | N |
| 5 | V149E | 14.76 | N |
| 7 | T26K | 14.63 | N |
| 8 | L7F | 14.54 | P |
| 9 | Q105F | 13.99 | N |
| 10 | Q105Y | 13.62 | N |
| 12 | Q105L | 12.31 | P |
| 19 | L7H | 11.63 | P |
| 22 | L7Y | 10.26 | P |
| 26 | T142Y | 9.67 | N |
| 27 | G30L | 9.62 | N |
| 30 | G30K | 8.69 | N |
| 32 | G30Q | 8.17 | N |
| 34 | L32F | 7.51 | P |
| 37 | T142K | 7.38 | N |
| 38 | F104E | 7.35 | N |
| 39 | F104Q | 7.32 | N |
| 44 | F104S | 7.17 | N |

## Concluding Discussion

Modeling protein mutation is crucial for protein design, protein evolution, and genetic disease analysis. The foundation for studying mutation is the ability to accurately model the side-chain conformations. In this research, we study the effect of protein mutation through the extent of predicted structural perturbation upon the mutation, specifically, through the extent of shift of the predicted side chains. To this end, we propose a computational method, namely OPUS-Mut. As shown in Figure 1, SI appendix, Fig. S1, SI appendix, Table S1 and SI appendix, Table S2, OPUS-Mut outperforms other backbone-dependent side-chain modeling methods including our previous method OPUS-Rota4 (19), either measured by all residues or measured by core residues exclusively. We evaluate the performance of OPUS-Mut on modeling mutation side chains. As shown in Figure 2, Figure 3, Figure 5 and SI appendix, Fig. S2, by comparing to experimental results, OPUS-Mut is capable of delivering satisfactory results for mutations.

OPUS-Mut can be used to infer the functional changes of the mutation. As exemplified in Figure 2f, from the side-chain prediction results of myoglobin mutant V68F, we can infer that the benzyl side chain, which partially fills the Xe4 cavity, may become a steric barrier to the ligand association. In addition, as shown in Figure 4, by comparing the differences of all side-chain dihedral angles (from χ_1_ to χ_4_) between the predicted wild-type side chains and the predicted mutant side chains, we find that the affected residues, whose predicted side chains are significantly shifted due to the mutation, can correspond well with the functional changes of the mutant. For example, the affected residues of the p53 Zn^2+^ region mutations are located at the Zinc-binding site (Figure 4a, Figure 4b, and Figure 4c), which indicate that the Zn^2+^ region may be affected in these mutants. In this case, OPUS-Mut can also be used to infer the affected residues caused by the mutation, therefore avoid the unwanted effect on specific sites in the case of protein engineering.

Although some mutations can lead to the perturbation of backbone, in our study, the backbone is set to be fixed as a first order approximation. Therefore, there could be some bias in our computation. However, we assume that the extent of the predicted side-chain shift can imply the interference of the corresponding mutation on the whole structure. From this point of view, the smaller differences (*S_diff_*) between the wild-type side chains and the mutant side chains, the smaller functional perturbation the mutation may cause. Therefore, using the side-chain modeling results with a fixed backbone as a first order approximation is a rational trade-off for studying protein point mutation.

OPUS-Mut can be used to identify the harmful (maximally disturbing) mutations. For a particular mutation, we sum up the differences (*S_diff_*) of all side-chain dihedral angles (from χ_1_ to χ_4_) between the predicted wild-type side chains and the predicted mutant side chains, and use it as an indicator for the extent of structural perturbation due to the mutation. By comparing with the experimentally determined functional changes data, the results show that larger structural perturbation corresponds to a higher probability of loss-of-function (Figure 6) or stability decrease (SI appendix, Fig. S4). Furthermore, as shown in Table 1 and Table 3, the harmful mutations are most likely in the top rank of the mutations with the maximal predicted structural perturbation.

OPUS-Mut can be used to identify the benign (minimally disturbing) mutations, which may guide us to construct a relatively low-homology mutant sequence but with similar structure. We screen all possible single site mutations for HIV-1 protease and T4 lysozyme. Among them, by preferably using the minimally disturbing mutations (with the lowest *S_diff_*), the constructed multiple mutation sequence tends to cause smaller structural perturbation with respect to wild-type structure, inferred from the predicted backbone generated by AlphaFold2 (Table 2 and SI appendix, Table S5). For example, the TM-score of the AlphaFold2 prediction for the HIV-protease mutant (SI appendix, Fig. S3) with 50% of mutation rate is 0.85, which is still reasonably high (the TM-score of the predicted wild-type structure is 0.93).

The 3D structure of protein determines its biological functions, therefore, studying protein mutation from structural perspective is important. There are two major types of methods for studying protein mutation from structural perspective. The first one is through protein structure prediction methods like AlphaFold2, which are capable of delivering all-atom structure for a particular sequence with relatively high accuracy based on multiple sequence alignment. However, their predictions are insensitive to the single site mutation, maybe due to the fact that the results of multiple sequence alignment are similar to each other when the difference between the wild-type and mutation sequences are relatively small. The second one is through backbone-dependent side-chain modeling methods, such as OPUS-Mut in this study. OPUS-Mut models the side chains using a fixed backbone (here, we use wild-type experimental backbone for both wild-type and mutation cases) as a first order approximation, and studies the effect of protein mutation through extent of predicted side-chain perturbation upon the mutation. OPUS-Mut adopts deep neural network to capture the local environment for each residue. Both kinds of method have their pros and cons, the former may be more suitable for mutations with larger mutation rate, and the latter may be more suitable for mutations with smaller mutation rate, even for single site mutation.

## Materials and methods

### Framework of OPUS-Mut

OPUS-Mut is mainly based on our previous work OPUS-Rota4 (19), but with some modifications. In calculation, only the protein residues are taken into consideration, other ligands are omitted. The neural network architecture of OPUS-Mut is shown in SI appendix, Fig. S5. In OPUS-Mut, we introduce the auxiliary loss *L_anglenorm_* from AlphaFold2 (11) to keep the predicted points close to the unit circle. For a dihedral angle *χ*, we output two predictions for sin (*χ*) and cos (*χ*), respectively. The loss function is listed as following:

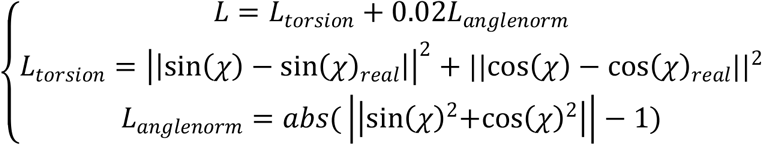

In addition, we add an additional node to predict the root-mean-square-deviation (RMSD) of the predicted side chain against its native counterpart for each residue. The side-chain RMSD ranges from 0 to 1, and segmented into 20 bins. The cross-entropy loss is used for the prediction.

OPUS-Mut is implemented in TensorFlow v2.4 (54) and trained on one NVIDIA Tesla V100. The Glorot uniform initializer and the Adam optimizer (55) are adopted. The initial learning rate is 1e-3 and it will be reduced by half when the accuracy of the validation set is decreased. The training process will stop after being reduced by four times. We train 5 models and the median of their outputs is used to make the final prediction.

### Datasets

In OPUS-Mut, the dataset used for training is the same as that in OPUS-Rota4 (19), which contains 10024 proteins in the training set and 983 proteins in the validation set, culled from the PISCES server on February 2017 (56). Note that none of the original structures and their related mutants mentioned in the four case studies of this paper are present in the dataset that is used to train the OPUS-Mut.

For evaluation, we use three native backbone test sets: CAMEO60, collected by OPUS-Rota3 (14), contains 60 hard targets released between January 2020 and July 2020 from the CAMEO website (57); CASP14, collected by OPUS-X (9), contains 15 FM targets downloaded from the CASP website (http://predictioncenter.org); CAMEO65, collected in this study, contains 65 hard targets released between May 2021 and October 2021 from the CAMEO website.

### Performance Metrics

To measure the accuracy of side-chain modeling methods, we use the percentage of correct prediction with a tolerance criterion 20° for all side-chain dihedral angles (from χ_1_ to χ_4_). The residue with more than 20 residues, between which the C_β_-C_β_ distance (C_α_ for Gly) is within 10 Å, is defined as core residue.

## Code Availability

The code and pre-trained models of OPUS-Mut as well as the datasets used in the study can be downloaded from http://github.com/OPUS-MaLab/opus_mut. They are freely available for academic usage only.

## Acknowledgements

The work was supported by Shanghai Municipal Science and Technology Major Project (No.2018SHZDZX01), and ZJLab. The work was also supported by National Key Research and Development Program of China (No. 2021YFF1200400).

## Reference

1. R. J. Kazlauskas, U. T. Bornscheuer, Finding better protein engineering strategies. Nat Chem Biol 5, 526–529 (2009).

2. A. N. Bullock, J. Henckel, A. R. Fersht, Quantitative analysis of residual folding and DNA binding in mutant p53 core domain: definition of mutant states for rescue in cancer therapy. Oncogene 19, 1245–1256 (2000).

3. B. Li, Y. T. Yang, J. A. Capra, M. B. Gerstein, Predicting changes in protein thermodynamic stability upon point mutation with deep 3D convolutional neural networks. Plos Comput Biol 16, e1008291 (2020).

4. D. E. Pires, D. B. Ascher, T. L. Blundell, mCSM: predicting the effects of mutations in proteins using graph-based signatures. Bioinformatics 30, 335–342 (2014).

5. M. Masso, Vaisman, II, Accurate prediction of enzyme mutant activity based on a multibody statistical potential. Bioinformatics 23, 3155–3161 (2007).

6. W. Torng, R. B. Altman, 3D deep convolutional neural networks for amino acid environment similarity analysis. Bmc Bioinformatics 18, 302 (2017).

7. R. Shroff et al., Discovery of Novel Gain-of-Function Mutations Guided by Structure-Based Deep Learning. Acs Synth Biol 9, 2927–2935 (2020).

8. T. Sanavia et al., Limitations and challenges in protein stability prediction upon genome variations: towards future applications in precision medicine. Comput Struct Biotechnol J 18, 1968–1979 (2020).

9. G. Xu, Q. Wang, J. Ma, OPUS-X: An Open-Source Toolkit for Protein Torsion Angles, Secondary Structure, Solvent Accessibility, Contact Map Predictions, and 3D Folding. Bioinformatics 38, 108–114 (2021).

10. J. Yang et al., Improved protein structure prediction using predicted interresidue orientations. Proc Natl Acad Sci U S A 117, 1496–1503 (2020).

11. J. Jumper et al., Highly accurate protein structure prediction with AlphaFold. Nature 596, 583–+ (2021).

12. M. Baek et al., Accurate prediction of protein structures and interactions using a three-track neural network. Science 373, 871–+ (2021).

13. G. R. Buel, K. J. Walters, Can AlphaFold2 predict the impact of missense mutations on structure? Nature Structural & Molecular Biology 29, 1–2 (2022).

14. G. Xu, Q. Wang, J. Ma, OPUS-Rota3: Improving Protein Side-Chain Modeling by Deep Neural Networks and Ensemble Methods. J Chem Inf Model 60, 6691–6697 (2020).

15. G. Xu, T. Ma, J. Du, Q. Wang, J. Ma, OPUS-Rota2: An Improved Fast and Accurate Side-Chain Modeling Method. J Chem Theory Comput 15, 5154–5160 (2019).

16. G. G. Krivov, M. V. Shapovalov, R. L. Dunbrack, Jr., Improved prediction of protein side-chain conformations with SCWRL4. Proteins 77, 778–795 (2009).

17. S. Liang, D. Zheng, C. Zhang, D. M. Standley, Fast and accurate prediction of protein side-chain conformations. Bioinformatics 27, 2913–2914 (2011).

18. M. Misiura, R. Shroff, R. Thyer, A. B. Kolomeisky, DLPacker: Deep Learning for Prediction of Amino Acid Side Chain Conformations in Proteins. bioRxiv 10.1101/2021.05.23.445347, 2021.2005.2023.445347 (2021).

19. G. Xu, Q. Wang, J. Ma, OPUS-Rota4: a gradient-based protein side-chain modeling framework assisted by deep learning-based predictors. Brief Bioinform 23, bbab529 (2022).

20. G. A. Ordway, D. J. Garry, Myoglobin: an essential hemoprotein in striated muscle. J Exp Biol 207, 3441–3446 (2004).

21. M. L. Quillin et al., Structural and functional effects of apolar mutations of the distal valine in myoglobin. J Mol Biol 245, 416–436 (1995).

22. K. Nienhaus, P. Deng, J. S. Olson, J. J. Warren, G. U. Nienhaus, Structural dynamics of myoglobin: ligand migration and binding in valine 68 mutants. J Biol Chem 278, 42532–42544 (2003).

23. M. L. Quillin, R. M. Arduini, J. S. Olson, G. N. Phillips, Jr., High-resolution crystal structures of distal histidine mutants of sperm whale myoglobin. J Mol Biol 234, 140–155 (1993).

24. S. Calhoun, V. Daggett, Structural Effects of the L145Q, V157F, and R282W Cancer-Associated Mutations in the p53 DNA-Binding Core Domain. Biochemistry-Us 50, 5345–5353 (2011).

25. A. Garufi et al., A fluorescent curcumin-based Zn(II)-complex reactivates mutant (R175H and R273H) p53 in cancer cells. J Exp clin Cancer Res 32, 72 (2013).

26. H. D. Shah, D. Saranath, V. Murthy, A molecular dynamics and docking study to screen anti-cancer compounds targeting mutated p53. J Biomol Struct Dyn 10.1080/07391102.2020.1839559, 1–10 (2020).

27. A. C. Joerger, H. C. Ang, A. R. Fersht, Structural basis for understanding oncogenic p53 mutations and designing rescue drugs. Proc Natl Acad Sci U S A 103, 15056–15061 (2006).

28. A. C. Joerger, H. C. Ang, D. B. Veprintsev, C. M. Blair, A. R. Fersht, Structures of p53 cancer mutants and mechanism of rescue by second-site suppressor mutations. J Biol Chem 280, 16030–16037 (2005).

29. A. C. Joerger, M. D. Allen, A. R. Fersht, Crystal structure of a superstable mutant of human p53 core domain - Insights into the mechanism of rescuing oncogenic mutations. Journal of Biological Chemistry 279, 1291–1296 (2004).

30. P. V. Nikolova, J. Henckel, D. P. Lane, A. R. Fersht, Semirational design of active tumor suppressor p53 DNA binding domain with enhanced stability. Proc Natl Acad Sci U S A 95, 14675–14680 (1998).

31. T. M. Rippin, S. M. Freund, D. B. Veprintsev, A. R. Fersht, Recognition of DNA by p53 core domain and location of intermolecular contacts of cooperative binding. J Mol Biol 319, 351–358 (2002).

32. Y. Cho, S. Gorina, P. D. Jeffrey, N. P. Pavletich, Crystal structure of a p53 tumor suppressor-DNA complex: understanding tumorigenic mutations. Science 265, 346–355 (1994).

33. V. Hornak, C. Simmerling, Targeting structural flexibility in HIV-1 protease inhibitor binding. Drug Discov Today 12, 132–138 (2007).

34. P. P. Mager, The active site of HIV-1 protease. Med Res Rev 21, 348–353 (2001).

35. P. J. Ala et al., Molecular basis of HIV-1 protease drug resistance: structural analysis of mutant proteases complexed with cyclic urea inhibitors. Biochemistry-Us 36, 1573–1580 (1997).

36. A. Ozen, K. H. Lin, N. Kurt Yilmaz, C. A. Schiffer, Structural basis and distal effects of Gag substrate coevolution in drug resistance to HIV-1 protease. Proc Natl Acad Sci U S A 111, 15993–15998 (2014).

37. M. Masso, Vaisman, II, Comprehensive mutagenesis of HIV-1 protease: a computational geometry approach. Biochem Biophys Res Commun 305, 322–326 (2003).

38. Z. Liu et al., Pulsed EPR characterization of HIV-1 protease conformational sampling and inhibitor-induced population shifts. Phys Chem Chem Phys 18, 5819–5831 (2016).

39. S. Pawar et al., Structural studies of antiviral inhibitor with HIV-1 protease bearing drug resistant substitutions of V32I, I47V and V82I. Biochem Biophys Res Commun 514, 974–978 (2019).

40. D. D. Loeb et al., Complete mutagenesis of the HIV-1 protease. Nature 340, 397–400 (1989).

41. R. Lapatto et al., X-ray analysis of HIV-1 proteinase at 2.7 A resolution confirms structural homology among retroviral enzymes. Nature 342, 299–302 (1989).

42. F. Sussman, M. C. Villaverde, A. Davis, Solvation effects are responsible for the reduced inhibitor affinity of some HIV-1 PR mutants. Protein Sci 6, 1024–1030 (1997).

43. N. M. King, M. Prabu-Jeyabalan, E. A. Nalivaika, C. A. Schiffer, Combating susceptibility to drug resistance: lessons from HIV-1 protease. Chem Biol 11, 1333–1338 (2004).

44. A. R. Poteete, L. W. Hardy, Genetic analysis of bacteriophage T4 lysozyme structure and function. J Bacteriol 176, 6783–6788 (1994).

45. L. H. Weaver, B. W. Matthews, Structure of bacteriophage T4 lysozyme refined at 1.7 Å resolution. Journal of Molecular Biology 193, 189–199 (1987).

46. J. A. Bell et al., Comparison of the crystal structure of bacteriophage T4 lysozyme at low, medium, and high ionic strengths. Proteins 10, 10–21 (1991).

47. S. Dao-pin, D. E. Anderson, W. A. Baase, F. W. Dahlquist, B. W. Matthews, Structural and thermodynamic consequences of burying a charged residue within the hydrophobic core of T4 lysozyme. Biochemistry-Us 30, 11521–11529 (1991).

48. D. W. Heinz, W. A. Baase, B. W. Matthews, Folding and function of a T4 lysozyme containing 10 consecutive alanines illustrate the redundancy of information in an amino acid sequence. Proc Natl Acad Sci U S A 89, 3751–3755 (1992).

49. J. Xu, W. A. Baase, E. Baldwin, B. W. Matthews, The response of T4 lysozyme to large-to-small substitutions within the core and its relation to the hydrophobic effect. Protein Sci 7, 158–177 (1998).

50. X. J. Zhang, B. W. Matthews, Conservation of solvent-binding sites in 10 crystal forms of T4 lysozyme. Protein Sci 3, 1031–1039 (1994).

51. M. Masso, T. Alsheddi, I. I. Vaisman, Accurate Prediction of Stability Changes in Bacteriophage T4 Lysozyme Upon Single Amino Acid Replacements. Ieee Int C Bioinform 10.1109/Bibm.2009.50, 26–30 (2009).

52. K. A. Bava, M. M. Gromiha, H. Uedaira, K. Kitajima, A. Sarai, ProTherm, version 4.0: thermodynamic database for proteins and mutants. Nucleic Acids Res 32, D120–121 (2004).

53. D. Rennell, S. E. Bouvier, L. W. Hardy, A. R. Poteete, Systematic mutation of bacteriophage T4 lysozyme. J Mol Biol 222, 67–88 (1991).

54. M. Abadi et al., TensorFlow: A system for large-scale machine learning. Proceedings of the 12th USENIX Symposium on Operating Systems Design and Implementation, 265–283 (2016).

55. D. P. Kingma, J. Ba, Adam: A Method for Stochastic Optimization. Proceedings of the 3rd International Conference on Learning Representations (2015).

56. J. Hanson, K. Paliwal, T. Litfin, Y. Yang, Y. Zhou, Improving prediction of protein secondary structure, backbone angles, solvent accessibility and contact numbers by using predicted contact maps and an ensemble of recurrent and residual convolutional neural networks. Bioinformatics 35, 2403–2410 (2019).

57. J. Haas et al., Continuous Automated Model EvaluatiOn (CAMEO) complementing the critical assessment of structure prediction in CASP12. Proteins 86 Suppl 1, 387–398 (2018).

