## Supplementary material for "OPUS-Mut: studying the effect of protein mutation through side-chain modeling": SI

**Fig. S1.** The percentage of correct prediction with a tolerance criterion 20° for all side-chain dihedral angles (from $\chi_1$ to $\chi_4$) of different methods measured by core residues.

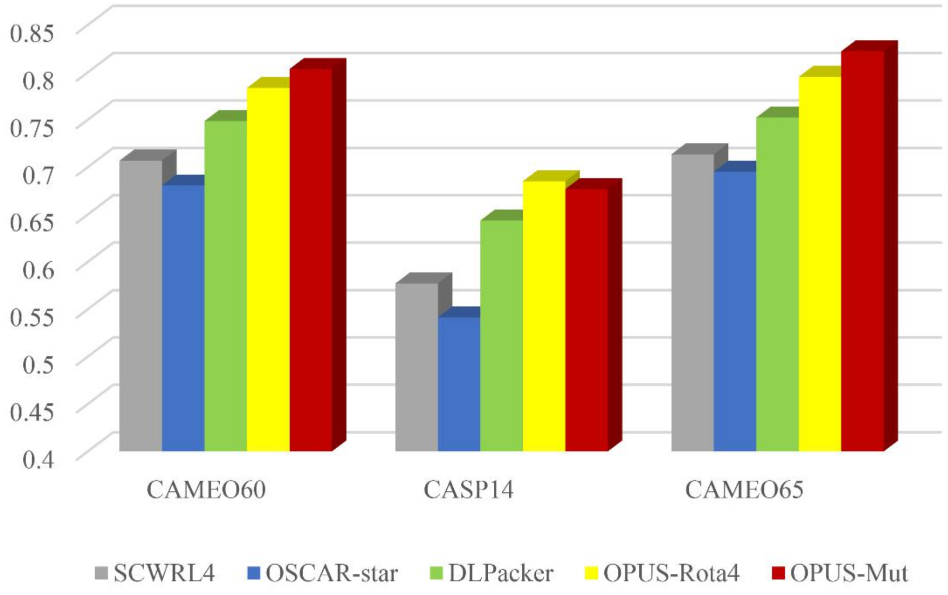

**Fig. S2.** Side-chain modeling results of *T*-p53C and its mutants. a) *T*-p53C (PDB: 1UOL); b) the Y220C mutant (PDB: 2J1X); c) the R282W mutant (PDB: 2J21); and d) the T123A/H168R/R249S mutant (PDB: 2BIQ). In all panels, the side chains from the crystal structures are marked in red for the mutation sites, and in blue for neighboring residues within 5 Å from the mutation sites. The side chains predicted by OPUS-Mut are marked in yellow.

.

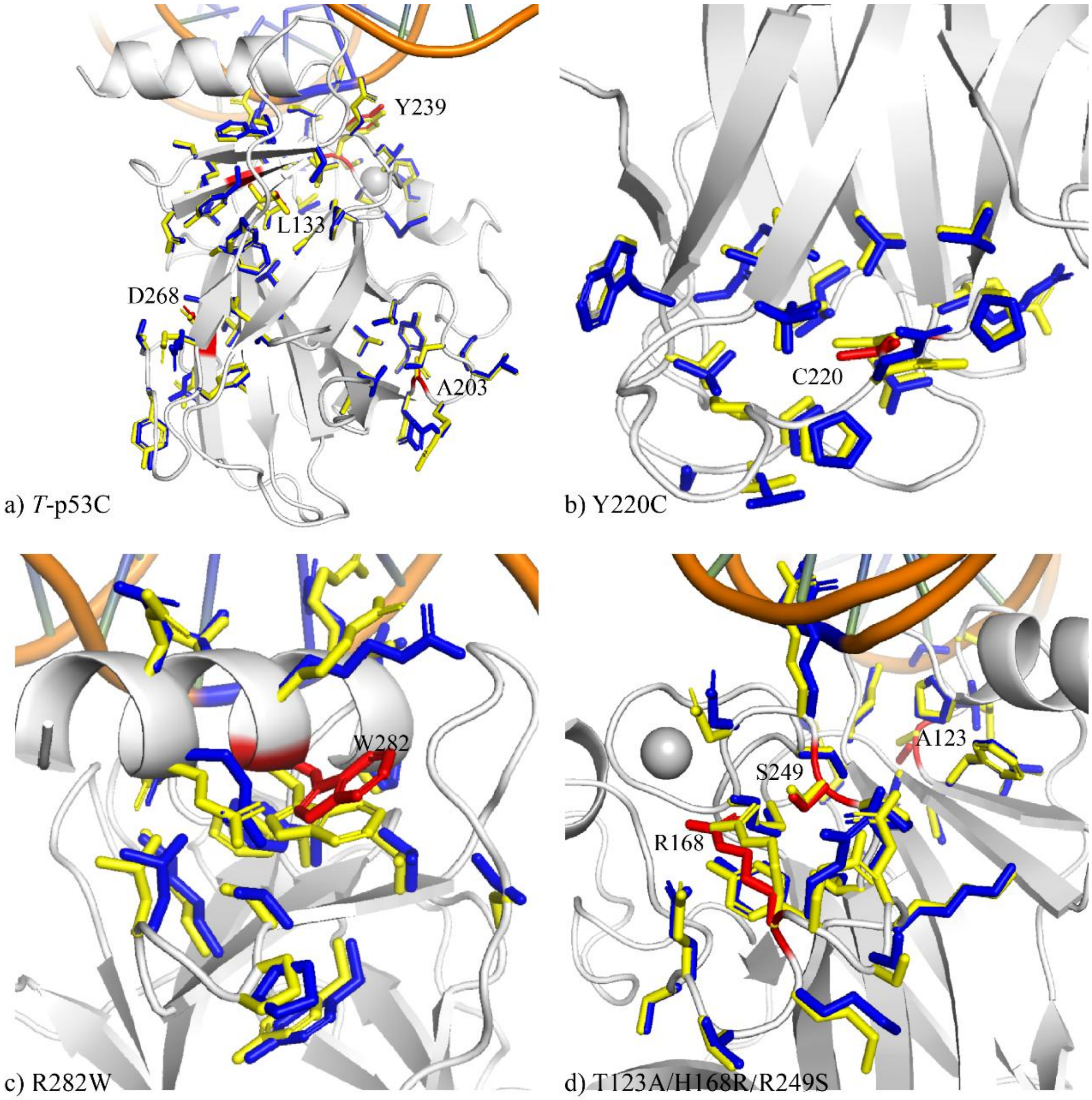

**Fig. S3.** The structure and sequence of the HIV-1 protease mutant with 50% of mutation rate. The blue structure is the wild-type crystal structure (PDB: 3PHV), the yellow structure is the mutant structure predicted by AlphaFold2. The mutated residues are marked in red on the mutant. The sequence above is the wild-type sequence, and the sequence below is the mutant sequence.

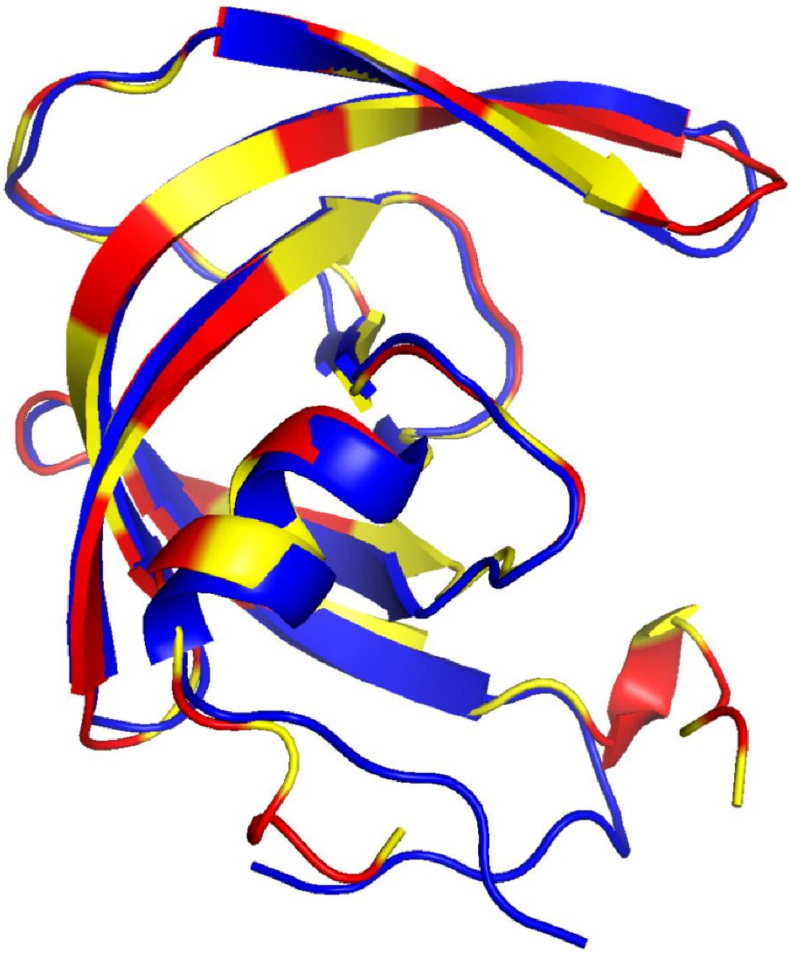

```
1        10        20        30        40        50        60
PQITLWQRPLVTIKIGGQLKEALLDTGADDTVLEEMSLPGRWKPKMIGGIGGFIKVRQYD
PEIALFHRPLVSIEIEAKEKSALLDTYANNTVLREMCLGGKWMGKQIGAHAADVYVEQYS

         70        80        90
QILIEICGHKAIGTVLVGPTPVNIIGRNLLTQIGCTLNF
EILIAVCANQPIASILVGKCHVNIIGKCLLTHIDCSMCF
```

**Fig. S4.** The summation of differences of all side-chain dihedral angles (from $\chi_1$ to $\chi_4$), either summed over all residues ($S_{diff}$) or summed over four binding site residues ($S_{diff\_critical}$), between the predicted wild-type side chains and the predicted mutant side chains on T4 lysozyme stability changes dataset. In the Box plot, "x" represents the mean of each group, the line inside the box represents the median of each group. The data from two groups (decreased stability (0), and increased stability group (1)) are shown. a) shows the differences ($S_{diff}$) summed over all residues. To avoid the influence of outliers, if the difference between the two residues is smaller than 1 degree, we set it to be 0; if it's smaller than 5, we set it to be 1; if it's smaller than 10, we set it to be 2; if it's smaller than 20, we set it to be 3; if it's larger than 20, we set it to be 4. b) shows the differences ($S_{diff\_critical}$) summed over four binding site residues (L32, F104, S117 and N132).

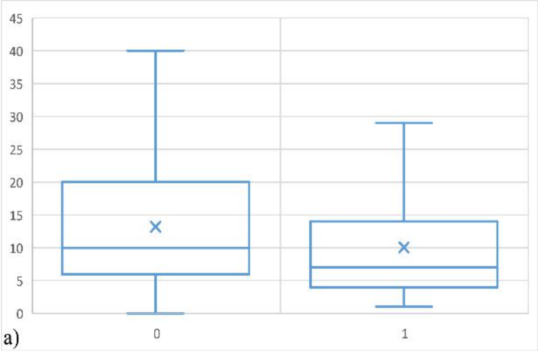


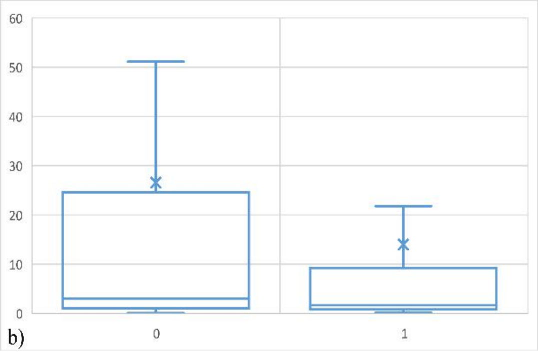

**Fig. S5.** The framework of OPUS-Mut. The trRosetta100 is a 100-d feature which is used to describe backbone distance ($C_\beta$-$C_\beta$) and orientation ($\omega$, $\theta_{ab}$, $\theta_{ba}$, $\varphi_{ab}$, $\varphi_{ba}$) contact information (1). The CSF15 refers to the relative CSF position (2) of the backbone atoms of a specific residue from its contact counterpart (3). The 2D features go through 32 dilated residual-convolutional blocks and an attention module, and output a 128-d vector for each residue. The 1D features are derived from protein backbone conformation. Beside the 41 features used in OPUS-Rota4 (3), an additional feature derived from DLPacker (4) is introduced to denote the side chain RMSD of prediction from its most similar conformation in the DLPacker's library. The 3DCNN is the side-chain density map for each residue, which uses the input from the DLPacker (OPUS) in OPUS-Rota4, it goes through a MPL unit and outputs a 512-d vector. The three outputs are combined and go through three modules which are identical to that in OPUS-TASS2 (5): modified Transformer module (6), Resnet module (7), and bidirectional Long-Short-Term-Memory module (8). All strides in the residual units are set to be one, and the batch size of OPUS-Mut is also set to be one.

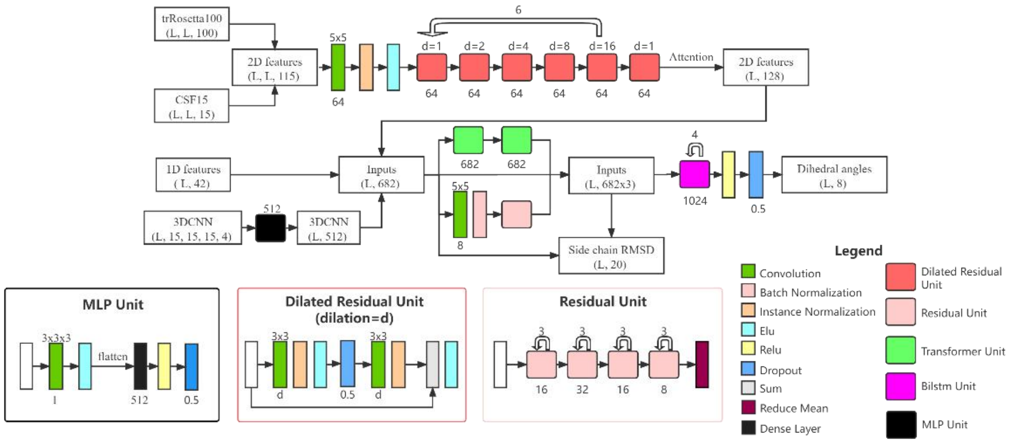

**Table S1**. The percentage of correct prediction of different residue types with a tolerance criterion 20° for all side-chain dihedral angles (from $\chi_1$ to $\chi_4$) of different methods on CAMEO65 measured by all residues. The best result for each type of residue is shown in boldface.

| | Total | SCWRL4 | OSCAR-star | DLPacker | OPUS-Rota4 | OPUS-Mut |
|---|---|---|---|---|---|---|
| ARG | 896 | 0.21 | 0.23 | 0.18 | 0.27 | **0.32** |
| ASN | 884 | 0.39 | 0.42 | 0.48 | 0.52 | **0.54** |
| ASP | 1066 | 0.48 | 0.50 | 0.58 | 0.60 | **0.62** |
| CYS | 264 | 0.78 | 0.81 | 0.84 | 0.84 | **0.87** |
| GLN | 668 | 0.22 | 0.23 | 0.24 | 0.32 | **0.34** |
| GLU | 1244 | 0.22 | 0.25 | 0.21 | 0.28 | **0.32** |
| HIS | 443 | 0.44 | 0.38 | 0.50 | 0.56 | **0.58** |
| ILE | 1217 | 0.70 | 0.70 | 0.68 | 0.73 | **0.75** |
| LEU | 1745 | 0.70 | 0.72 | 0.76 | 0.77 | **0.79** |
| LYS | 1065 | 0.19 | 0.21 | 0.10 | 0.21 | **0.21** |
| MET | 398 | 0.34 | 0.36 | 0.33 | 0.41 | **0.44** |
| PHE | 741 | 0.72 | 0.70 | 0.81 | 0.76 | **0.85** |
| PRO | 833 | **0.78** | 0.76 | 0.78 | 0.74 | 0.78 |
| SER | 1144 | 0.56 | 0.62 | 0.67 | **0.70** | 0.69 |
| THR | 1027 | 0.79 | 0.81 | 0.81 | **0.83** | 0.83 |
| TRP | 297 | 0.71 | 0.64 | 0.84 | **0.90** | 0.90 |
| TYR | 652 | 0.75 | 0.71 | 0.86 | 0.79 | **0.88** |
| VAL | 1174 | 0.83 | 0.84 | 0.87 | 0.89 | **0.89** |

**Table S2**. The percentage of correct prediction of different residue types with a tolerance criterion 20° for all side-chain dihedral angles (from $\chi_1$ to $\chi_4$) of different methods on CAMEO65 measured by core residues. The best result for each type of residue is shown in boldface.

| | Total | SCWRL4 | OSCAR-star | DLPacker | OPUS-Rota4 | OPUS-Mut |
|---|---|---|---|---|---|---|
| ARG | 138 | 0.36 | 0.33 | 0.30 | 0.44 | **0.48** |
| ASN | 176 | 0.50 | 0.53 | 0.61 | 0.69 | **0.70** |
| ASP | 145 | 0.59 | 0.57 | 0.74 | 0.78 | **0.79** |
| CYS | 145 | 0.79 | 0.79 | 0.85 | 0.86 | **0.88** |
| GLN | 136 | 0.32 | 0.29 | 0.35 | 0.54 | **0.57** |
| GLU | 125 | 0.37 | 0.34 | 0.30 | 0.42 | **0.49** |
| HIS | 119 | 0.54 | 0.39 | 0.63 | 0.74 | **0.75** |
| ILE | 605 | 0.78 | 0.77 | 0.76 | 0.82 | **0.85** |
| LEU | 853 | 0.79 | 0.77 | 0.84 | 0.86 | **0.87** |
| LYS | 102 | 0.37 | 0.46 | 0.23 | 0.43 | **0.49** |
| MET | 187 | 0.47 | 0.43 | 0.42 | 0.55 | **0.57** |
| PHE | 350 | 0.79 | 0.73 | 0.87 | 0.85 | **0.93** |
| PRO | 178 | **0.85** | 0.82 | 0.84 | 0.81 | 0.84 |
| SER | 322 | 0.65 | 0.70 | 0.77 | 0.81 | **0.82** |
| THR | 268 | 0.86 | 0.87 | 0.88 | 0.90 | **0.90** |
| TRP | 132 | 0.81 | 0.70 | 0.87 | 0.95 | **0.95** |
| TYR | 270 | 0.81 | 0.72 | 0.93 | 0.84 | **0.95** |
| VAL | 574 | 0.86 | 0.86 | 0.89 | 0.92 | **0.93** |

**Table S3**. The HIV-1 protease mutagenesis dataset (0, 1, and 2 denote negative, intermediate, and positive phenotypes, respectively). The mutations and their phenotypes are separated by space character in the table.

| | | | | | | |
|---|---|---|---|---|---|---|
| P1S 1 | E21Q 2 | L33N 0 | G51A 2 | I66M 1 | P81R 0 | C95Y 0 |
| P1L 2 | E21V 2 | L33D 0 | G51K 0 | I66V 2 | V82I 2 | C95R 0 |
| P1H 2 | E21K 2 | L33Y 0 | G51V 0 | I66S 0 | V82L 2 | T96P 0 |
| Q2E 2 | A22G 1 | L33E 0 | G51C 0 | I66T 0 | V82A 1 | T96S 1 |
| I3L 2 | A22T 0 | E34A 2 | G52A 0 | I66F 2 | V82G 0 | T96R 1 |
| I3T 1 | A22D 0 | E35S 2 | G52S 0 | C67G 2 | V82T 2 | L97S 0 |
| I3S 0 | L23V 1 | E35A 1 | G52V 0 | C67F 2 | V82F 1 | L97F 0 |
| I3N 2 | L23P 0 | E35L 1 | G52D 0 | C67W 1 | V82D 0 | N98D 1 |
| T4S 2 | L23Q 0 | E35K 1 | F53Y 2 | C67Y 2 | N83D 0 | N98S 2 |
| T4R 1 | L23R 0 | E35M 1 | F53I 2 | G68V 0 | N83K 0 | N98Y 1 |
| L5V 0 | L24M 1 | E35G 1 | F53L 2 | G68E 0 | N83I 0 | F99A 0 |
| L5H 0 | L24V 1 | M36R 0 | F53V 2 | H69R 2 | N83T 0 | F99P 0 |
| W6G 2 | L24S 0 | S37G 2 | I54L 2 | H69L 2 | I84L 1 | F99S 0 |
| W6L 2 | L24F 0 | S37R 2 | I54S 1 | H69Y 2 | I84M 0 | F99L 2 |
| W6C 2 | D25A 0 | L38V 1 | I54T 1 | H69D 0 | I84K 0 | F99D 0 |
| Q7P 0 | D25Y 0 | L38F 1 | I54F 1 | H69N 1 | I85L 1 | F99K 0 |
| Q7H 2 | D25H 0 | P39A 1 | K55R 1 | H69Q 2 | I85S 0 | F99R 0 |
| R8G 0 | T26A 0 | P39T 2 | K55T 2 | K70T 2 | I85F 0 | F99Y 1 |
| R8Q 1 | T26I 0 | P39L 1 | K55N 2 | K70N 2 | I85N 0 | F99E 0 |
| P9S 2 | T26K 0 | P39R 1 | K55Q 2 | A71P 0 | G86S 0 | |
| P9T 0 | T26R 0 | G40E 0 | V56I 0 | A71S 1 | G86I 0 | |
| P9H 0 | G27A 0 | G40R 0 | V56L 0 | A71L 2 | G86E 0 | |
| P9R 0 | G27P 0 | R41K 2 | V56G 0 | I72T 2 | G86N 0 | |
| L10V 2 | G27I 0 | W42G 2 | R57K 2 | I72L 2 | G86Q 0 | |
| L10H 1 | G27V 0 | W42R 0 | R57G 0 | I72V 2 | G86R 0 | |
| L10R 1 | G27R 0 | K43N 2 | R57S 0 | G73R 0 | G86K 0 | |
| V11A 1 | G27D 0 | K43Q 2 | R57T 0 | G73Q 0 | R87S 0 | |
| V11G 0 | A28S 0 | P44T 2 | R57I 0 | G73T 1 | N88A 1 | |
| T12S 2 | A28G 0 | P44R 2 | Q58E 2 | G73C 2 | L89V 2 | |
| I13V 2 | A28T 0 | K45R 2 | Q58P 0 | G73V 1 | L90V 0 | |
| I13L 1 | A28E 0 | K45T 2 | Q58K 1 | G73A 1 | L90S 0 | |
| I13M 1 | D29E 0 | K45Q 2 | Q58H 1 | G73K 0 | L90F 0 | |
| K14T 2 | D29N 0 | M46L 2 | Y59D 0 | G73P 0 | L90W 0 | |
| K14M 2 | D29A 0 | I47M 0 | Y59C 0 | T74S 2 | T91A 2 | |
| K14E 1 | D29G 0 | I47S 0 | D60E 2 | T74I 0 | T91P 0 | |
| K14N 2 | D29V 0 | I47T 0 | D60V 1 | V75E 0 | T91N 2 | |
| K14Q 2 | D30E 2 | I47D 0 | Q61P 1 | V75G 0 | Q92P 0 | |
| I15V 2 | D30N 1 | I47E 0 | Q61L 1 | L76I 0 | Q92L 2 | |
| I15T 0 | D30Y 1 | I47R 0 | Q61R 2 | V77G 0 | Q92H 1 | |
| I15R 0 | T31S 1 | G48S 2 | I62L 2 | V77E 0 | I93M 2 | |
| G16V 1 | T31A 0 | G48T 1 | L63I 2 | G78A 0 | I93V 2 | |
| G16W 1 | T31I 0 | G48E 1 | L63V 2 | G78V 0 | I93T 1 | |

| | | | | | |
|---|---|---|---|---|---|
| G17W 1 | T31R 0 | G48H 2 | L63P 2 | G78E 0 | I93F 2 |
| G17E 2 | V32L 2 | G49V 0 | L63R 2 | P79A 2 | I93N 0 |
| Q18L 2 | V32E 0 | G49R 0 | I64L 2 | P79S 2 | G94A 0 |
| Q18H 2 | L33V 2 | I50L 2 | I64T 1 | P79T 1 | G94V 0 |
| Q18R 2 | L33A 0 | I50M 1 | I64R 0 | T80P 0 | G94R 1 |
| L19T 2 | L33G 0 | I50T 0 | E65V 0 | P81A 0 | C95G 0 |
| K20M 2 | L33S 0 | I50N 0 | E65K 0 | P81T 0 | C95S 0 |
| K20Q 1 | L33T 0 | I50R 0 | I66L 2 | P81H 0 | C95F 1 |

**Table S4**. The T4 lysozyme stability changes dataset. 0 and 1 denote decrease and increase the stability respectively. The mutations and their stability changes are separated by space character in the table.

| | | | | | | |
|---|---|---|---|---|---|---|
| I3A 0 | L39A 0 | I58A 0 | S90H 0 | M106L 1 | V131A 1 | F153I 0 |
| I3C 1 | N40A 1 | I58T 0 | L91A 0 | E108V 1 | V131D 1 | F153L 1 |
| I3D 0 | N40D 1 | I58Y 0 | L91M 0 | T109D 1 | V131E 1 | F153M 0 |
| I3E 0 | A41D 1 | T59A 0 | L91P 0 | T109N 1 | V131G 0 | F153V 0 |
| I3F 0 | A41S 0 | T59D 0 | D92N 0 | V111A 0 | V131I 1 | R154E 1 |
| I3G 0 | A41V 1 | T59G 0 | A93P 1 | V111F 0 | V131L 1 | G156D 0 |
| I3L 1 | A42F 0 | T59N 0 | A93S 0 | V111I 0 | V131M 1 | T157A 0 |
| I3M 0 | A42G 0 | T59S 0 | A93T 1 | G113A 1 | V131S 0 | T157C 0 |
| I3P 0 | A42I 0 | T59V 0 | V94A 0 | G113E 1 | V131T 0 | T157D 0 |
| I3S 0 | A42K 0 | K60H 0 | R96C 0 | T115A 0 | N132F 1 | T157E 0 |
| I3T 0 | A42L 0 | K60P 1 | R96H 0 | T115E 1 | N132I 1 | T157F 0 |
| I3V 0 | A42S 0 | L66A 0 | A98C 0 | N116A 1 | N132M 1 | T157G 0 |
| I3W 0 | A42V 0 | L66P 0 | A98F 0 | N116D 1 | L133A 0 | T157H 0 |
| I3Y 0 | K43A 0 | F67A 0 | A98I 0 | S117A 1 | L133D 0 | T157I 0 |
| M6A 0 | S44A 1 | N68A 0 | A98L 0 | S117F 1 | L133F 0 | T157L 0 |
| M6I 0 | S44C 0 | Q69P 0 | A98M 0 | S117I 1 | L133M 0 | T157N 0 |
| M6L 0 | S44D 0 | V71A 0 | A98S 0 | S117V 1 | A134S 0 | T157R 1 |
| L7A 0 | S44E 1 | D72P 0 | A98T 0 | L118A 0 | K135C 0 | T157S 0 |
| E11A 1 | S44F 1 | A73S 0 | A98V 0 | L118I 0 | K135E 0 | T157V 0 |
| E11F 1 | S44G 0 | A74P 0 | A98W 0 | L118M 0 | W138Y 0 | D159C 1 |
| E11H 1 | S44H 1 | V75T 0 | L99A 0 | R119A 0 | T142C 0 | A160T 0 |
| E11M 1 | S44I 1 | G77A 1 | L99F 0 | R119C 0 | P143A 0 | N163D 0 |
| E11N 0 | S44K 1 | I78A 0 | L99G 0 | R119E 1 | N144D 1 | |
| R14K 0 | S44L 1 | I78M 0 | L99I 0 | R119H 0 | N144E 1 | |
| K16E 1 | S44M 1 | I78V 0 | L99M 0 | R119M 1 | N144H 1 | |
| I17A 0 | S44N 0 | L79C 0 | L99V 0 | M120A 0 | A146C 0 | |
| D20A 0 | S44P 0 | R80K 0 | I100A 0 | M120K 0 | A146I 0 | |
| D20N 1 | S44Q 1 | A82P 1 | I100M 0 | M120L 1 | A146T 0 | |
| D20S 1 | S44R 1 | A82S 0 | I100V 0 | M120Y 0 | A146V 0 | |
| D20T 1 | S44T 1 | K83H 0 | N101A 0 | L121A 0 | K147E 1 | |
| E22K 1 | S44V 1 | L84A 0 | M102K 0 | L121M 0 | V149A 0 | |
| Y25G 0 | S44W 1 | L84M 0 | M102L 0 | Q122A 0 | V149C 0 | |
| T26Q 0 | S44Y 1 | P86A 0 | M102T 0 | Q123A 0 | V149G 0 | |
| T26S 1 | E45A 1 | P86C 0 | M102V 0 | Q123E 1 | V149I 0 | |
| I27A 0 | L46A 0 | P86D 1 | V103A 0 | K124G 1 | V149S 0 | |
| G28A 0 | D47A 0 | P86G 0 | V103I 0 | W126R 0 | V149T 0 | |
| I29A 0 | K48A 0 | P86H 0 | V103M 0 | E128A 1 | I150V 0 | |
| G30A 1 | A49S 0 | P86I 0 | F104A 0 | E128K 0 | T151S 1 | |
| G30F 0 | I50A 0 | P86L 0 | Q105A 0 | A129F 0 | T152A 0 | |
| H31N 0 | G51D 0 | P86R 0 | Q105E 0 | A129L 0 | T152C 0 | |
| L33A 0 | C54T 1 | P86S 0 | Q105G 0 | A129M 0 | T152I 0 | |
| P37A 1 | C54V 0 | P86T 0 | Q105M 0 | A129V 0 | T152S 0 | |

| | | | | | |
|---|---|---|---|---|---|
| S38A 0 | C54Y 0 | V87A 0 | M106A 0 | A129W 0 | T152V 1 |
| S38D 1 | N55C 0 | V87I 0 | M106I 1 | A130G 0 | F153A 0 |
| S38N 0 | N55G 0 | V87T 0 | M106K 0 | A130S 0 | F153C 0 |

**Table S5**. The TM-score of each T4 lysozyme (with 164 residues in length) multiple mutation sequence predicted by AlphaFold2. "Type 1" represents the TM-score of the sequences with 10%, 20%, 30%, 40%, and 50% of mutation rate, constructed by preferably using the minimally disturbing mutations with the lowest $S_{diff}$. "Type 2" represents the TM-score of the multiple mutation sequences constructed by preferably using the maximally disturbing mutations with the highest $S_{diff}$.

| | Wild-type | 10% | 20% | 30% | 40% | 50% |
|---|---|---|---|---|---|---|
| Type 1 | 0.99 | 0.98 | 0.98 | 0.97 | 0.95 | 0.84 |
| Type 2 | - | 0.82 | 0.84 | 0.80 | 0.69 | 0.55 |

### SI Reference


1. J. Yang *et al.*, Improved protein structure prediction using predicted interresidue orientations. *Proc Natl Acad Sci U S A* **117**, 1496-1503 (2020).
2. G. Xu, T. Q. Ma, T. W. Zang, Q. H. Wang, J. P. Ma, OPUS-CSF: A C-atom-based scoring function for ranking protein structural models. *Protein Sci* **27**, 286-292 (2018).
3. G. Xu, Q. Wang, J. Ma, OPUS-Rota4: a gradient-based protein side-chain modeling framework assisted by deep learning-based predictors. *Briefings in Bioinformatics* 10.1093/bib/bbab529, bbab529 (2021).
4. M. Misiura, R. Shroff, R. Thyer, A. B. Kolomeisky, DLPacker: Deep Learning for Prediction of Amino Acid Side Chain Conformations in Proteins. *bioRxiv* 10.1101/2021.05.23.445347, 2021.2005.2023.445347 (2021).
5. G. Xu, Q. Wang, J. Ma, OPUS-X: an open-source toolkit for protein torsion angles, secondary structure, solvent accessibility, contact map predictions and 3D folding. *Bioinformatics* **38**, 108-114 (2022).
6. G. Xu, Q. H. Wang, J. P. Ma, OPUS-TASS: a protein backbone torsion angles and secondary structure predictor based on ensemble neural networks. *Bioinformatics* **36**, 5021-5026 (2020).
7. K. M. He, X. Y. Zhang, S. Q. Ren, J. Sun, Deep Residual Learning for Image Recognition. *Proc Cvpr Ieee* 10.1109/Cvpr.2016.90, 770-778 (2016).
8. S. Hochreiter, J. Schmidhuber, Long short-term memory. *Neural Comput* **9**, 1735-1780 (1997).
